# N-acetylaspartate improves cell survival when glucose is limiting

**DOI:** 10.1101/2020.05.28.114629

**Authors:** H. Furkan Alkan, Katharina E. Walter, Hubert Hackl, Matthew G. Vander Heiden, Tobias Madl, Juliane G. Bogner-Strauss

## Abstract

N-acetylasparate (NAA), previously considered a brain-specific metabolite, is found in several cancers. However, whether it plays a role in tumor growth or survival is not fully understood. We provide evidence that NAA prevents cell death in low-glucose conditions via sustaining intracellular UDP-N-acetylglucosamine (UDP-GlcNac) levels, suppressing endoplasmic reticulum (ER) stress, and enabling continued protein synthesis. NAA production is critical for *in vivo* tumor growth where lower glucose levels are present than those in cell culture. Furthermore, the breakdown of NAA leads to ER stress and cell death, suggesting that the role of NAA in low-glucose is independent of its catabolism to produce aspartate or acetate. Together, these data suggest NAA can support the growth of some tumors by helping them cope with glucose limitations *in vivo*.

**Highlights:** Endogenous N-acetylaspartate (NAA) production boosts tumor growth

NAA supports cell survival in low glucose via suppressing ER stress

Breaking down NAA limits tumor growth and induces ER stress *in vivo*

The role of NAA to rescue low glucose is independent of donating acetate or aspartate

**In brief:** Cancer cells need N-acetylaspartate to avoid ER stress and cell death when glucose availability is low.

## Introduction

N-acetylaspartate (NAA) is an amino acid derivative abundantly found in the brain (Moffett et al., 2013). Although it is commonly associated with brain health, there is no consensus on the precise role of NAA within the central nervous system (CNS). Because low NAA levels in the brain are associated with hypomyelination, one role of NAA is suggested to be supplying acetate for oligodendrocytes. The proposed model includes the synthesis of NAA in the neurons that express Aspartate-N-acetyltransferase (NAT8L) enzyme and shuttling of NAA to the surrounding oligodendrocytes where it is cleaved by the aspartoacylase (ASPA) enzyme (Moffett et al., 2013). The loss of ASPA leads to Canavan disease, characterized by the accumulation of NAA and hypomyelination in the brain. On the other hand, ASPA and NAT8L double knockout mice display improved myelination and brain development compared to single ASPA knockouts (Sohn et al., 2017), conflicting with the hypothesis that the major role of NAA is providing acetate to oligodendrocytes.

Contrary to initial assumptions that NAA is a brain-specific metabolite, NAA production has also been observed outside the CNS, particularly in adipocytes (Bogner-Strauss, 2017). Intracellular NAA levels boost browning of white adipocytes and increase lipolysis and mitochondrial uncoupling in brown adipocytes. Furthermore, NAA can regulate de novo lipogenesis and autophagy through altering cytosolic acetyl-CoA levels in immortalized brown adipocyte cells (Pessentheiner et al., 2013; Huber et al., 2019). In addition to adipocytes, NAA levels are also detected in plasma and in tumor biopsies from lung adenocarcinoma patients (Lou et al., 2015). NAT8L expression was also associated with increased tumor growth and poor survival in ovarian cancer (Zand et al., 2016). Despite accumulating reports on increased NAT8L expression and NAA levels in several solid tumors, the mechanistic role of NAA remains enigmatic.

The tumor microenvironment can introduce several metabolic limitations for cell proliferation, including nutrient starvation and hypoxia, that are absent in standard cell culture conditions (Muir et al., 2018). In this study, we revealed that increased Nat8l expression leads to slower cell proliferation *in vitro* while boosting tumor growth *in vivo*. We identified that this phenotype is due to Nat8l-overexpressing cells surviving and proliferating better in low-glucose environments. Mechanistically, we showed that NAA increases uridine diphosphate-N-acetylglucosamine (UDP-GlcNac) biosynthesis, which improves cell survival and proliferation via sustaining protein synthesis and suppressing ER stress following glucose starvation (Palorini et al., 2013; de la Cadena et al., 2014; Wang et al., 2014). These findings argue that a major role of NAA in cancers is to limit ER stress and promote survival when glucose is limiting.

## Results

### Nat8l overexpression reduces proliferation *in vitro* while increasing tumor growth *in vivo*

NAA is an amino acid derivative produced from L-aspartate and acetyl-CoA by the NAT8L enzyme and can be broken down into L-aspartate and acetate by ASPA (Figure 1A). Interestingly, patient survival data from The Cancer Genome Atlas (TCGA) shows that regulation of NAT8L and ASPA genes favoring increased NAA levels (high NAT8L or low ASPA) are often correlated with poor prognosis (Figure S1A). Given that high NAT8L expression was associated with low ASPA expression within the same type of cancer, we hypothesized that the role of NAA in cancers might be independent of its catabolism. Therefore, we manipulated the gene expressions of both Nat8l and Aspa to investigate the role of NAA in cancer cell proliferation. Because endogenous NAA levels were not detected in mouse Lewis lung carcinoma (LLC1) cells using gas chromatography-mass spectrometry (GC-MS) (Alkan et al., 2018), we first overexpressed Nat8l in these cells to investigate the impact of increased NAA production. As expected, Nat8l overexpression boosts intracellular NAA levels without having any effect on Aspa expression (Figure 1B, 1C, S1B). Interestingly, Nat8l overexpressing LLC1 cells (Nat8l o/e) proliferate slower than control (PuroN) cells in full media conditions (Figure 1D). On the other hand, Nat8l overexpression improves the growth of allograft LLC1 tumors (Figure 1E, Figure S1B), suggesting that the role of NAA in cancer could be environment-dependent.

**Figure 1.**
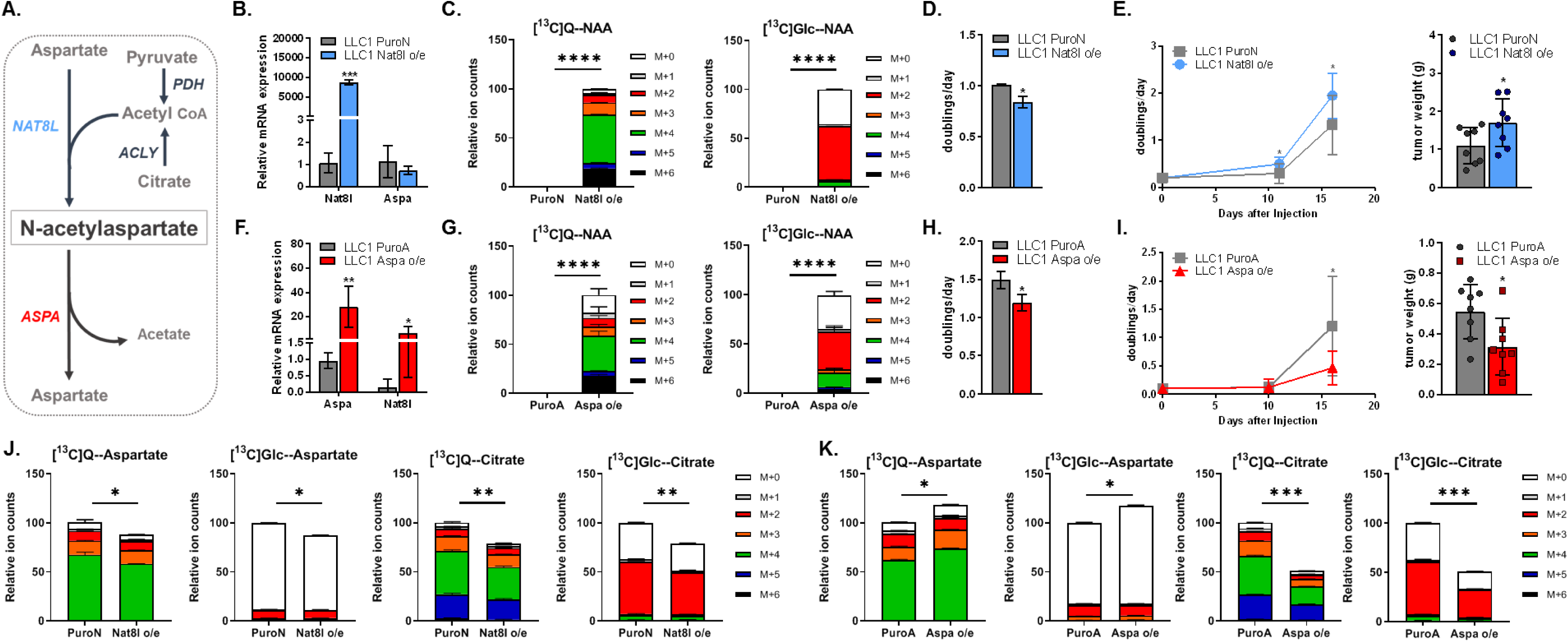
Impacts of Nat8l and Aspa overexpressions on cell proliferation, tumor growth and TCA cycle metabolism. **A)** Schematic demonstration of enzymes and metabolic intermediates involved in N-acetylaspartate synthesis (Nat8l) and breakdown (Aspa). Pyruvate dehydrogenase (PDH) and ATP-citrate lyase (ACLY) are also shown as alternative sources of Acetyl-CoA. **B, F)** Verification of **B)** Nat8l and **F)** Aspa overexpressions in LLC1 cells using qPCR analysis. **C, G)** Relative levels of N-acetylaspartate (NAA) in control (PuroN and PuroA), **C)** Nat8l o/e and **G)** Aspa o/e LLC1 cells. 24 hours tracing of [U^13^C] glutamine (left) or [U^13^C]glucose (right) into NAA is also shown. **D, H)** Proliferation rate (shown as doublings/day) of LLC1 control (PuroN and PuroA), **D)** Nat8l o/e and **H)** Aspa o/e cells cultured in DMEM (n=3). **E, I)** (left) Tumor volume and (right) tumor weights of control (PuroN and PuroA), **E)** Nat8l o/e and **I)** Aspa o/e LLC1 allografts 11 and 16 days after the injections into the flanks of C56BL/6 mice (n=8). **J, K)** Relative intracellular levels of aspartate and citrate in control (PuroN and PuroA), **J)** Nat8l o/e and **K)** Aspa o/e LLC1 cells cultured in DMEM without pyruvate (n=3). 24 hours tracing of [U^13^C] glutamine (Q) or [U^13^C]glucose (Glc) into aspartate and citrate are also shown. All figures denote mean ± SDs. Significance levels: * p ≤ 0.05, ** p ≤ 0.01, *** p ≤ 0.001, **** p ≤ 0.00001

Next, we overexpressed the NAA-cleaving enzyme Aspa in LLC1 cells (Figure 1A, 1F, S1C). Of note, Aspa overexpressing cells (Aspa o/e) were challenging to recover after retroviral infection; however, those Aspa o/e cells that were recovered displayed prominent upregulation of Nat8l (Figure 1F), implying that even though endogenous levels of NAA and Nat8l in LLC1 are below the detection limit, some intracellular NAA could still be important for these cells. In addition, the Nat8l upregulation in Aspa overexpressing (Aspa o/e) cells results in higher NAA levels than parental cells *in vitro* (Figure 1G), despite increased Aspa expression. Next, we measured the proliferation rate of Aspa o/e cells and observed that Aspa overexpression leads to slower proliferation rate *in vitro*, similar to Nat8l overexpression (Figure 1H). However, Aspa overexpression does not phenocopy Nat8l overexpression *in vivo* and yields smaller tumors (Figure 1I, Figure S1C). Nevertheless, it is important to note that intracellular NAA levels in Nat8l o/e cells were about 50 times higher than those in Aspa o/e cells (Figure S1D), and NAA levels were 10-times higher in Nat8l o/e tumors compared to controls while Aspa overexpression did not alter NAA levels *in vivo* (Figure S1E, S1F). Of note, mice bearing Nat8l overexpressing tumors also have elevated NAA levels in their plasma, suggesting that tumors also secrete NAA (Figure S1G).

Because Aspa o/e cells have a perpetual cycle of NAA synthesis and catabolism, this may disrupt metabolism via diverting aspartate and Acetyl-CoA from other essential pathways that could overweigh any beneficial role of increased NAA levels. Interestingly, Aspa o/e LLC1 cells also lost Aspa expression several weeks after selection and culture *in vitro*. Because Aspa o/e cells had difficulties recovering from puromycin selection, we regenerated the cells using a vector with hygromycin (hygro) resistance (Figure S1H). We found that Aspa expression in hygro-selected Aspa o/e cells also slowly reduced over time without significant changes in Nat8l expression, further suggesting that high Aspa expression might be harmful to proliferating cells (Figure S1H, S1I).

Because intracellular aspartate levels could be limiting for proliferation (Birsoy et al., 2015; Sullivan et al., 2015; Gui et al., 2016; Alkan and Bogner-Strauss, 2019) we hypothesized that increased NAA production may deplete aspartate in cells, which might explain the reduced proliferation rate of Nat8l o/e cells *in vitro*. Interestingly, despite high NAA levels, intracellular aspartate concentrations were minimally changed in both Nat8l o/e and Aspa o/e cells compared to control (Figure 1J, 1K), suggesting that NAA production is not limited by aspartate availability in culture. Interestingly, Aspa o/e cells have slightly increased intracellular aspartate despite slower proliferation (Figure 1K). Consistently, transient overexpression of Aspa in Nat8l o/e cells further reduces the proliferation (Figure S1J), indicating that slower proliferation of Nat8l o/e cells is independent of aspartate levels. On the other hand, levels of citrate, one presumed source of acetate for cytosolic NAA (Huber et al., 2019, Jiang et al., 2017), were reduced in both Aspa and Nat8l overexpressing cells (Figure 1J, 1K). In addition, steady-state labeling with [U^13^C]-Glucose and [U^13^C]-Glutamine revealed that the major source of the acetate moiety of NAA is glucose, while glutamine provides the aspartate carbons (Figure 1C, 1G, 1J, 1K). Interestingly, the labeling pattern of NAA was not identical between Aspa and Nat8l overexpressing cells, implying that endogenously upregulated Nat8l might have a different subcellular localization than the constitutively overexpressed Nat8l. Of note, proportional glucose incorporation to citrate was slightly increased in Nat8l o/e cells and reduced in Aspa o/e cells (Figure 1J, 1K), suggesting that a shift in glucose metabolism might have occurred in these cells contributing to slower proliferation *in vitro*.

### NAA improves cell survival when glucose is limiting

Many nutrients are less abundant in the tumor microenvironment compared to cell culture (Muir et al., 2017). To test whether Nat8l overexpressing cells adapt the limited resources better than control cells, we systematically removed the major carbon sources; glucose, glutamine, and pyruvate from the media and measured the proliferation rate. Strikingly, Nat8l overexpressing cells proliferate better than the control cells in glucose-depleted media while growing slower in full media conditions (Figure 2A). On the other hand, Aspa overexpressing cells do not display any advantage in glucose starvation (Figure 2A). Interestingly, providing exogenous NAA improves the proliferation and survival of both control and Nat8l o/e cells following glucose starvation (Figure 2B). Next, we inhibited glucose metabolism using 2-deoxyglucose (2-DG) when glucose is present. Consistent with the previous observation, Nat8l o/e cells proliferate better following 2-DG treatment, and exogenous NAA again improves cell viability in both control and Nat8l o/e cells (Figure 2B). Remarkably, Aspa o/e cells proliferate poorly in all conditions and NAA supplementation recovers Aspa o/e cells minimally compared to controls (Figure 2B), consistent with the hypothesis that breakdown of NAA might have adverse effects on cells.

**Figure 2.**
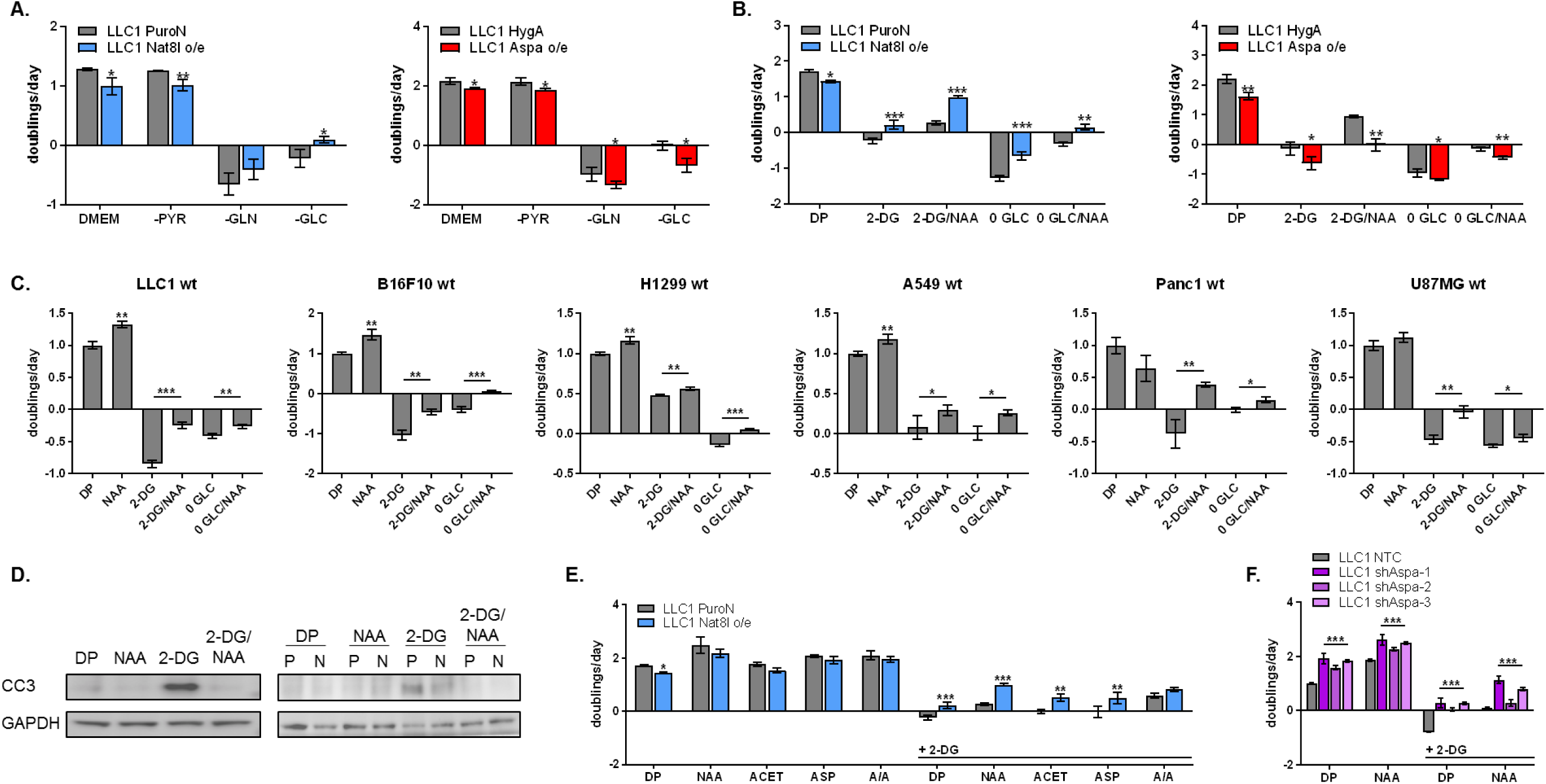
NAA improves cell proliferation and survival in low glucose. **A)** Proliferation/survival rate (shown as doublings/day) of LLC1 controls (PuroN, HygA), Nat8l o/e and Aspa o/e cells cultured in full media (DMEM), DMEM without pyruvate (-PYR), DMEM without glucose and pyruvate (-GLC), and DMEM without glutamine and pyruvate (-GLN) for 48 hours (n=3). All conditions contain 10% dialysed FBS. **B)** Proliferation/survival rate (shown as doublings/day) of LLC1 controls (PuroN, HygA), Nat8l o/e and Aspa o/e cells in DMEM without pyruvate (DP) in the absence or presence of 5 mM 2-deoxyglucose (2-DG), 10 mM N-acetylaspartate (NAA) or without glucose (0 GLC) for 72 (Nat8l o/e) or 48 (Aspa o/e) hours (n=3). All conditions contain 10% dialysed FBS. **C)** Proliferation/survival rate (shown as doublings/day) of wild-type LLC1, B16F10, H1299, A549, PANC1 and U87MG cells in DMEM without pyruvate (DP) in the absence or presence of 10 mM 2-deoxyglucose (2-DG), 10 mM N-acetylaspartate (NAA) or without glucose (0 GLC) for 72 hours (n=3). All conditions contain 10% dialysed FBS. **D)** Western Blot analysis of Cleaved Caspase 3 (CC3) protein expression in wild-type (left), and Nat8l overexpression (right), in the presence and absence of 5 mM 2-deoxyglucose (2-DG) and 10 mM N-acetylaspartate (NAA). **E)** Proliferation/survival rate (shown as doublings/day) of LLC1 controls (PuroN) and Nat8l o/e cells in DMEM without pyruvate (DP) in the absence or presence of 5 mM 2-deoxyglucose (2-DG), 10 mM N-acetylaspartate (NAA), 10 mM Aspartate (ASP), 10 mM Acetate (ACET), or the combination of aspartate and acetate (A/A) for 48 hours, as indicated (n=3). All conditions contain 10% dialysed FBS. **F)** Proliferation/survival rate (shown as doublings/day) of LLC1 control (NTC) and Aspa-KD (shAspa-1, shAspa-2, and shAspa-3) cells in DMEM without pyruvate (DP) in the absence or presence of 5 mM 2-deoxyglucose (2-DG), 10 mM N-acetylaspartate (NAA) for 48 hours, as indicated (n=3). All conditions contain 10% dialysed FBS. All figures denote mean ± SDs. Significance levels: * p ≤ 0.05, ** p ≤ 0.01, *** p ≤ 0.001

To test whether NAA supplementation rescues glucose starvation in other systems, we examined wild-type LLC1, B16F10, A549, H1299, PANC1, and U87MG cells. Strikingly, NAA treatment improved the proliferation/survival of all the cells considered following either glucose depletion or 2-DG treatment, although the impact of either glucose limitation or NAA supplementation varied among cell lines (Figure 2C). Consistently, Nat8l overexpressing A549 and H1299 cells survived better following glucose starvation or 2-DG treatments (Figure S2A-D). Interestingly, exogenous NAA improves the proliferation of melanoma (B16F10) and lung cancer (LLC1, A549, and H1299) cells even in full media conditions while having unfavorable or no effect on pancreas (PANC1) and glioblastoma (U87MG) cell lines, respectively. This implies some tissue-specific need for NAA. Furthermore, we measured the levels of cleaved-caspase 3 (CC3), a marker of apoptosis, in cells treated with 2-DG. As expected, 2-DG treatment increases CC3 levels in wild-type LLC1 cells (Figure 2D). Consistent with the cell counts, Nat8l o/e cells, but not Aspa o/e cells, have reduced CC3 levels upon 2-DG treatment compared to controls (Figure 2D, S2E). Strikingly, 2-DG-induced CC3 is almost completely abolished upon NAA supplementation (Figure 2D), indicating that NAA prevents apoptosis. Altogether, these findings suggest that both exogenous and endogenous NAA can improve cell proliferation and survival when glucose metabolism is inhibited.

Next, we investigated the impact of aspartate and acetate (two components of NAA) in 2-DG treated cells. Interestingly, both acetate and aspartate improve the survival of 2-DG treated LLC1 cells (Figure 2D). However, the ability of NAA, or the dual treatment with aspartate and acetate, to rescue survival are superior compared to aspartate or acetate supplementation alone, implying that aspartate and acetate might contribute to NAA production to have an effect (Figure 2D). We then knocked-down Aspa (Aspa-KD) in LLC1 cells to determine whether blocking the breakdown of NAA would impair its ability to rescue cell viability in glucose limitation (Figure S2F). Interestingly, Aspa-KD cells proliferate faster in regular media compared to controls (Figure 2F). In addition, Aspa-KD cells display increased survival upon 2-DG treatment. Importantly, NAA supplementation also improves the cell number of Aspa-KD cells with 2-DG treatment, suggesting that the cell-autonomous breakdown of NAA is not necessary to rescue glucose limitation.

### NAA improves ER stress response and protein synthesis rate

To investigate why Nat8l overexpressing cells display improved survival in low glucose environment or upon 2-DG treatment, we characterized their metabolic phenotype using a non-targeted ^1^H-NMR analysis. Interestingly, one of the significantly increased metabolite in Nat8l overexpressing LLC1 cells (apart from NAA itself) that could be related to surviving glucose starvation is UDP-N-acetylglucosamine (UDP-GlcNac) (Figure 3A). Moreover, UDP-glucose levels are also higher in Nat8l o/e LLC1 cell *in vitro*, while UDP-galactose levels are not significantly changed (Figure 3B, 3C). On the other hand, either NAA supplementation or Nat8l overexpression do not regulate hexoseamine biosynthesis pathway on a transcription level (Figure S3). Glucose depletion leads to impaired hexosamine biosynthesis and protein glycosylation which activates the unfolded protein response (Palorini et al., 2013). Even though we cannot directly link NAA to UDP-GlcNac metabolism using known biochemical pathways, we hypothesized that Nat8l o/e cells might endure glucose limitation-induced ER stress better than controls due to having increased intracellular UDP-GlcNac levels (Figure 3D).

**Figure 3.**
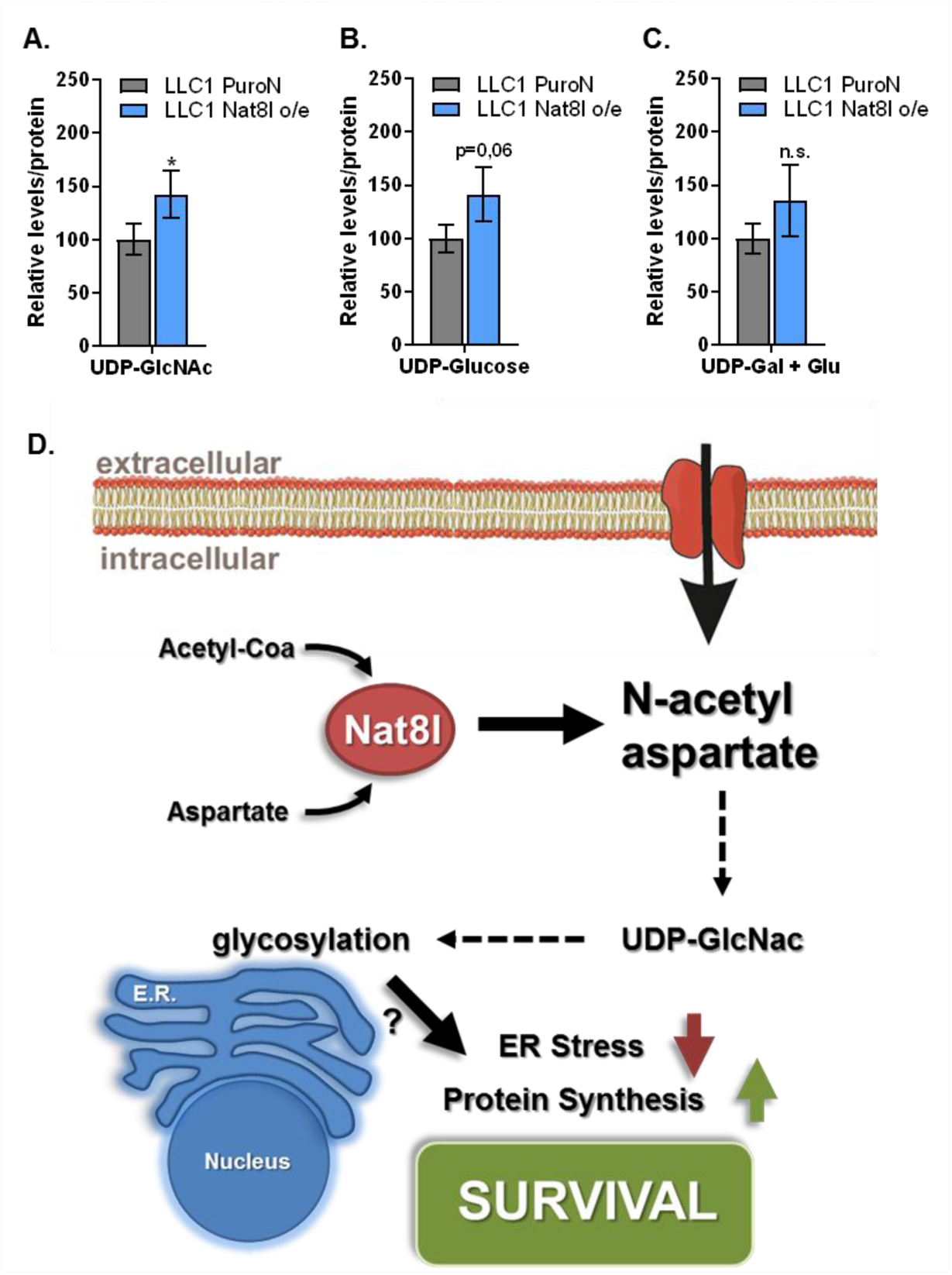
NAA increases intracellular levels of UDP-sugars. **(A-C)** Relative levels of intracellular (A) UDP-N-acetylglucosamine (UDP-GlcNac) (B) UDP-glucose and (C) UDP-galactose + UDP-glucuronate mixture (UDP-Gal + Glu) in control (PuroN) and Nat8l overexpressing (Nat8l o/e) LLC1 cells, determined by NMR spectroscopy (n=3). Mean ± SDs. are shown. Significance levels: * p ≤ 0.05 **(D)** Schematic showing of potential impact of N-acetylaspartate for cell survival in low glucose conditions. UDP-GlcNac sustains protein glycosylation which reduces ER stress and promotes protein synthesis, leading to improved cell survival.

To determine whether Nat8l overexpressing cells have an improved ER stress response, we measured gene expressions of various ER stress markers in the presence and absence of 2-DG. As expected, 2-DG treatment strongly induced the mRNA expressions of C/EBP homologous protein (CHOP), activating transcription factor 6 (Atf6), spliced X-box binding protein (Xbp1s), and 90kDa heat shock protein (Hsp90b1) (Figure 4A). Remarkably, this induction was significantly weaker in Nat8l o/e cells, suggesting that increased levels of intracellular UDP-GlcNac might suppress ER stress in these cells (Figure 4A). Furthermore, exogenous NAA treatment also improves the ER stress response in control and wild-type LLC1 cells (Figure 4A, B). Aspa o/e cells display mild levels of ER stress even without 2-DG treatment, which is further increased in the presence of 2-DG (Figure 4C). In Aspa o/e cells, NAA treatment did not improve ER stress response, consistent with the hypothesis that NAA does not function as a source for aspartate or acetate to improve cell survival upon glucose limitations (Figure 4C). In addition, CHOP and BiP protein expressions are also upregulated upon 2-DG treatment and recovered by Nat8l overexpression or NAA supplementation (Figure S4A).

**Figure 4.**
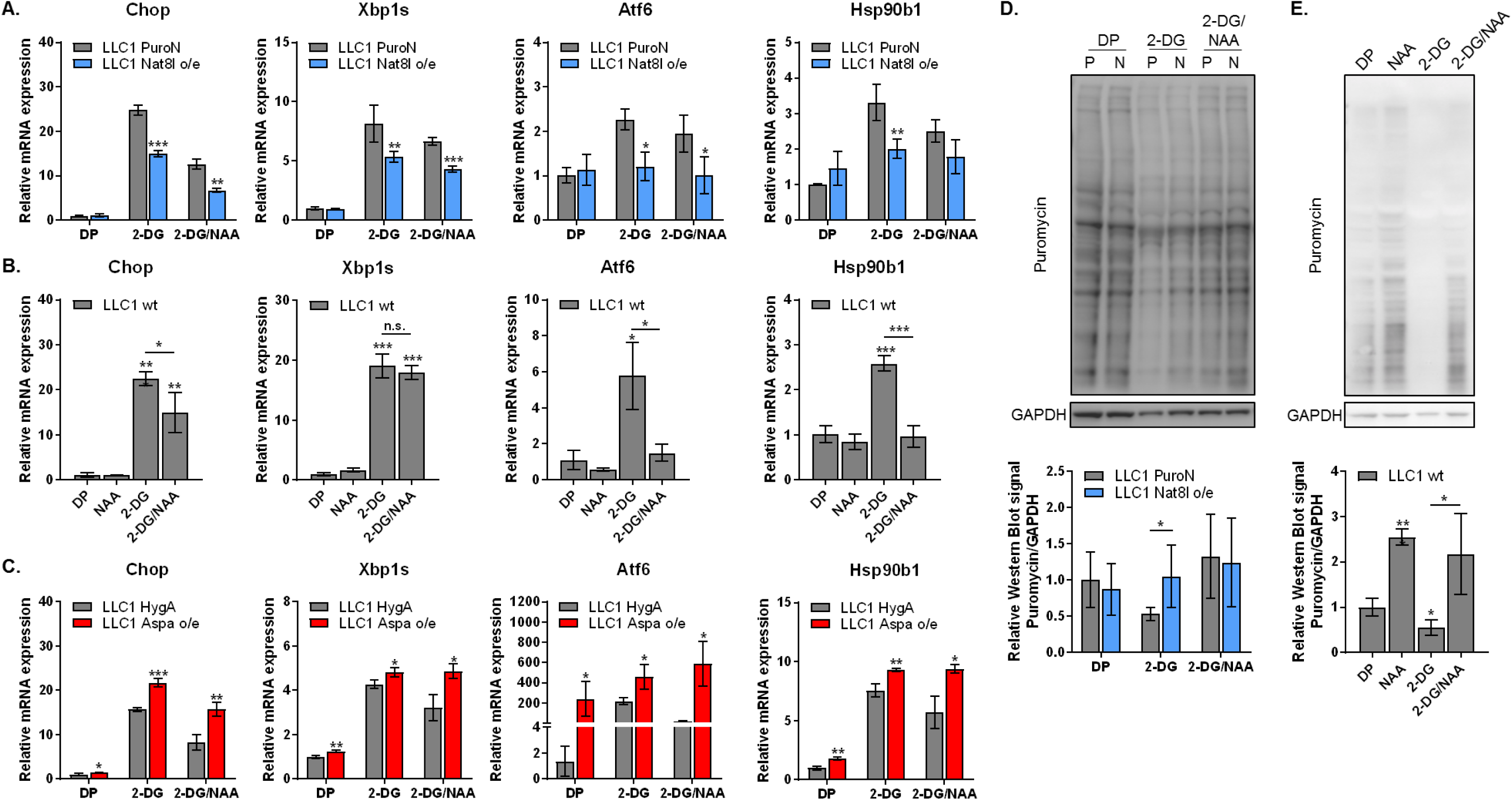
NAA improves ER stress response and protein synthesis when glucose metabolism is inhibited. **A-C)** Relative mRNA levels of Chop, Xbp1s, Atf6, and Hsp90b1 in **A)** control (PuroN) and Nat8l o/e, **B)** wild-type and **C)** control (HygA) and Aspa o/e LLC1 cells in the presence and absence of 5 mM 2-deoxyglucose (2-DG) and 10 mM N-acetylaspartate (NAA). Rplp0 is used as reference gene (n=3). **D, E)** Western blot analysis of the puromycin incorporation into protein synthesis after 15 minutes of 90 µM puromycin treatment in the presence and absence of 5 mM 2-deoxyglucose (2-DG) and 10 mM N-acetylaspartate (NAA). All figures denote mean ± SDs. Significance levels: * p ≤ 0.05, ** p ≤ 0.01, *** p ≤ 0.001

Next, we investigated the impact of NAA supplementation on 2-DG-induced ER stress in A549 and H1299 cells. Consistent with our findings from LLC1 cells, 2-DG treatment leads to elevated markers of ER stress in these human lung cancer cell lines which was diminished by exogenous NAA (Figure S4B). We also tested whether Nat8l o/e cells have an improved ER stress signature *in vivo*, however, Nat8l o/e LLC1 tumors did not exhibit any difference in mRNA expressions of ER stress-related genes compared to controls (Figure S4C). This might be due to Nat8l o/e tumors growing faster than the controls (Figure 1E). On the other hand, Aspa o/e LLC1 tumors had higher baseline expressions of ER stress marker genes *in vivo*, as observed *in vitro* (Figure S4D, 4C).

Because UDP-GlcNac reduces protein misfolding (Denzel et al., 2014; Vincenz and Hartl, 2014; Wang et al., 2014), we hypothesized that Nat8l overexpression or NAA supplementation might also improve protein synthesis. To determine the protein synthesis rate, we treated the cells with puromycin for a short time and traced its incorporation into proteins using western blot. As expected, 2-DG treatment reduces puromycin incorporation in LLC1 cells (Figure 4D, 4E). Importantly, Nat8l o/e cells exhibit higher puromycin incorporation upon 2-DG treatment (Figure 4D). Furthermore, NAA supplementation also increases the protein synthesis rate in wild-type LLC1, A549, and H1299 cells even without 2-DG treatment (Figure 4E, S4E). These findings suggest that one major role of NAA is to modulate ER stress and protein synthesis particularly when glucose is limiting.

## Discussion

These findings argue that NAA improves cancer cell proliferation and survival when glucose metabolism is compromised. These data are consistent with previous reports suggesting Nat8l expression is upregulated in certain cancers and is beneficial for proliferation (Lou et al., 2015; Zand et al., 2016). Mechanistically, we show that NAA is a mediator of ER stress and protein synthesis through boosting intracellular UDP-GlcNac levels, which is crucial for cell survival when glucose is absent.

Intratumor levels of some nutrients, including glucose, can be lower than those in standard cell culture conditions (Sullivan et al., 2019). Therefore, the ability to survive this stress *in vivo* could be more crucial than how fast a cancer cell can proliferate when every nutrient is present. Because Nat8l overexpressing cells endure glucose starvation better than the control cells, this might explain the inconsistency in proliferation rate between *in vitro* and *in vivo* experiments. Because both Nat8l o/e and Aspa-KD cells can cope with glucose starvation better than the control cells, and NAA supplementation can still improve survival/proliferation in Aspa-KD cells, we speculate that it is not the breakdown products of NAA (aspartate or acetate), but rather NAA itself is needed to improve ER stress and protein synthesis rate when glucose is limiting.

One consequence of glucose starvation is decreased UDP-GlcNAc levels (Palorini et al., 2013; de la Cadena et al., 2014). Disruption in hexosamine biosynthesis causes unfolded protein accumulation that can lead to ER stress, and eventually apoptosis if adaptive mechanisms fail (Denzel at al., 2014; Vincenz and Hartl, 2014; Wang et al., 2014). We speculate that Nat8l overexpressing cells or cells treated with NAA survive better in low-glucose conditions because NAA increases intracellular UDP–GlcNac levels. Because NAA-breakdown into acetate is not required for rescuing glucose limitations, we speculate that NAA does not compensate for the ATP-generating function of glycolysis but rather decreases ER stress via another unknown mechanism.

Aspartate is one of the substrates needed for pyrimidine biosynthesis (Alkan et al., 2018; Sullivan et al., 2015) therefore UDP-GlcNac levels could be affected by aspartate bioavailability. However, without glucose, the ribose backbone of UDP is still absent which cannot be rescued by aspartate, unless through gluconeogenesis. In addition, Aspa o/e cells have higher aspartate levels while doing poorly in low glucose and NAA supplementation can rescue survival of Aspa-knockdown cells, suggesting that aspartate moiety of N-acetylaspartate is not limiting in glucose starvation.

Despite the strong evidence that NAA levels improve ER stress-response and cell survival, through which mechanism UDP levels are increased in Nat8l o/e cells is unclear. We suspect that this does not occur through regulating transcription because we did not observe changes in expression of genes involved in hexosamine biosynthesis in Nat8l o/e cells (Wang et al., 2014) (Figure S3). Also, this effect is likely to be independent of a non-specific activity of the Nat8l enzyme as exogenous NAA supplementation phenocopies the impact of Nat8l overexpression in low-glucose conditions. Therefore, we speculate that the effect of NAA could be either post-translational via acetylation of certain enzymes, histones or DNA, or a metabolic effect independent from just supplying aspartate and acetate. Nevertheless, further studies are needed to unravel a potential role of NAA in signaling or altering other aspects of metabolism.

These findings provide evidence that one major role of NAA in cancer metabolism is to prevent ER stress and apoptosis when glucose is limiting, suggesting that NAA levels could be a predictor of the outcomes of some metabolism-based therapies. In addition, this study might provide insight into the role of NAA in the central nervous system. Further studies are required to understand whether NAA has similar anti-apoptotic roles in the brain by preventing ER stress.

### Limitations of the study

Endogenous NAA levels in wild-type LLC1 cells were close to limit of detection; however, despite very low Nat8l expression in these cells (Ct values ≥ 33), NAA still correlated with phenotypes observed in Aspa o/e and Aspa-KD cells. In addition, we could not find an antibody to detect endogenous Nat8l and Aspa protein in any of the cell lines tested, although Aspa mRNA expressions were much higher than Nat8l. One of the major limitations of our study is that Aspa overexpressing cells strongly upregulate Nat8l expression and result in higher intracellular NAA levels. Moreover, Aspa overexpression in LLC1 cells was unstable over time. Finally, despite repeated attempts, Aspa overexpression could not be obtained in the human cells considered, further suggesting toxicity of this enzyme for some cancer cells.

## Supplemental Information

Supplemental Information includes 4 Supplemental Figures.

## AUTHOR CONTRIBUTIONS

H.F.A designed and performed *in vivo* tumor growth, metabolite extraction, GC-MS, and a part of the proliferation experiments. H.F.A and K.E.W designed and K.E.W performed qPCR, western blot, and a part of cell proliferation/survival experiments. H.H. conducted bioinformatics analysis. T.M performed NMR measurements and carried out NMR data analysis. M.G.V.H provided substantial guidance and shared lab space and equipment for GC-MS experiments. H.F.A and J.G.B-S constructed the study and H.F.A., K.E.W., and J.G.B-S wrote the manuscript.

## ACKNOWLEDGEMENTS

This work was funded by the Austrian Science Fund FWF SFB LIPTOX F3018, P27108, and W1226 DK “Metabolic and Cardiovascular Disease”. M.G.V.H. acknowledges the support from the Lustgarten Foundation, SU2C, the Ludwig Center at MIT, the MIT Center for Precision Cancer Medicine, and the NCI (P30 CA1405141, R01 CA168653). M.G.V.H. is also an HHMI faculty scholar. H.F.A was supported by Austrian Marshall Plan Scholarship. We acknowledge the support of NAWI Graz and the technical support of Thomas Schreiner, Wolfgang Krispel, Anna Springer, Sarah Stryeck, and other members of Bogner-Strauss lab. T.M. was supported by Austrian Science Foundation Grants P28854, I3792, and DK-MCD W1226; Austrian Research Promotion Agency (FFG) Grants 864690 and 870454; the Integrative Metabolism Research Center Graz; Austrian Infrastructure Program 2016/2017, the Styrian Government (Zukunftsfonds), and BioTechMed-Graz (Flagship project).

## MATERIALS & METHODS

### KEY RESOURCES TABLE

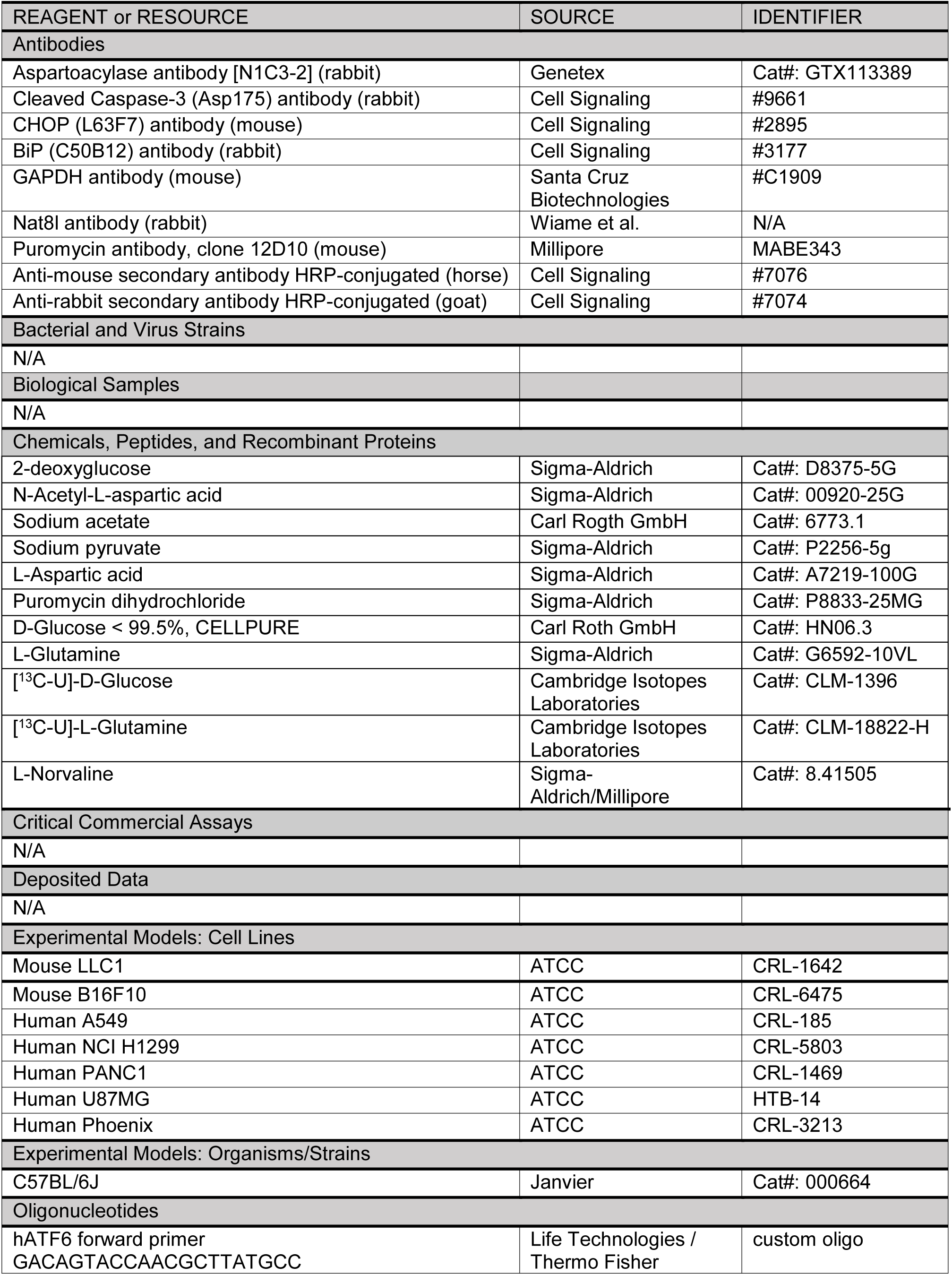
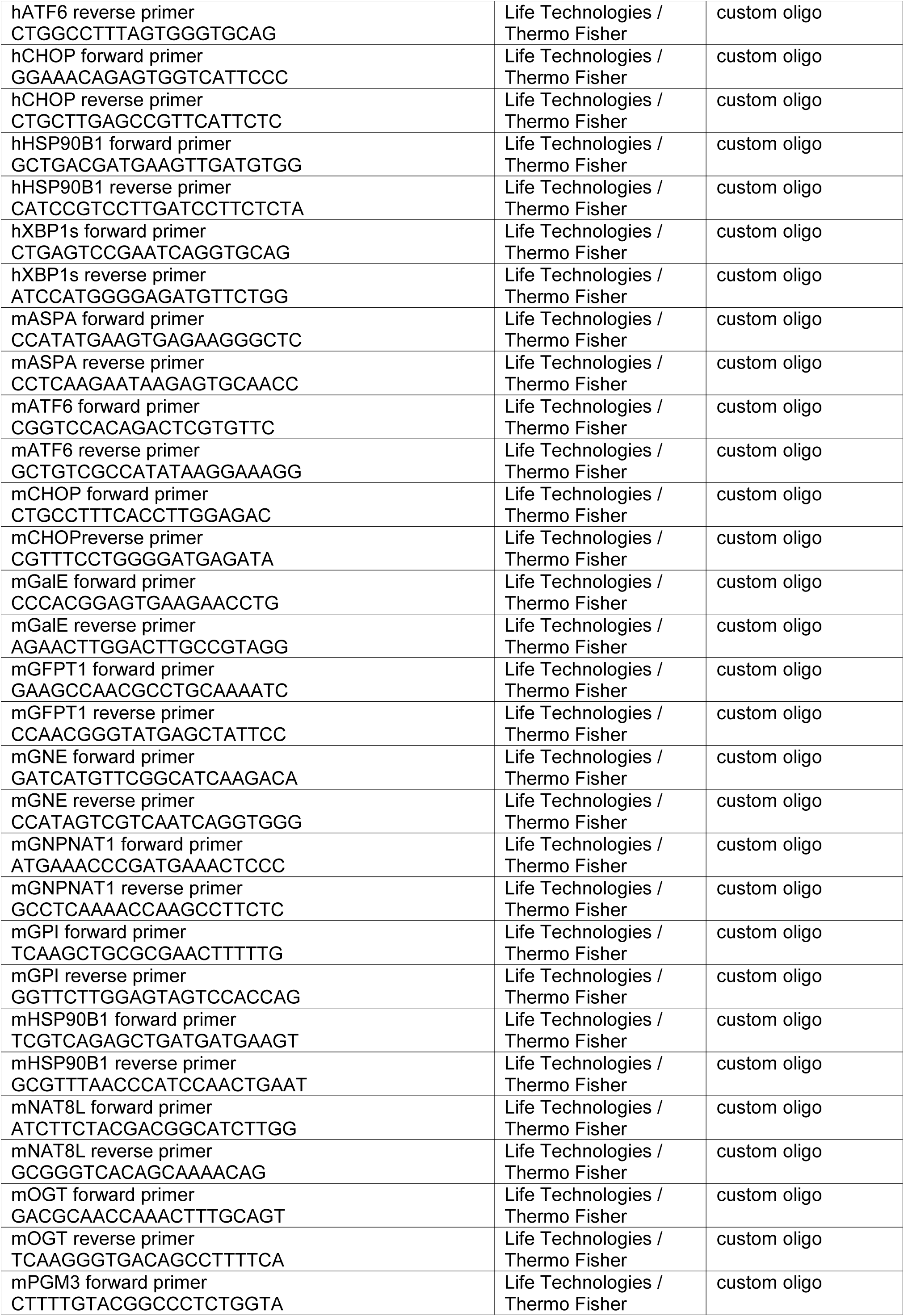
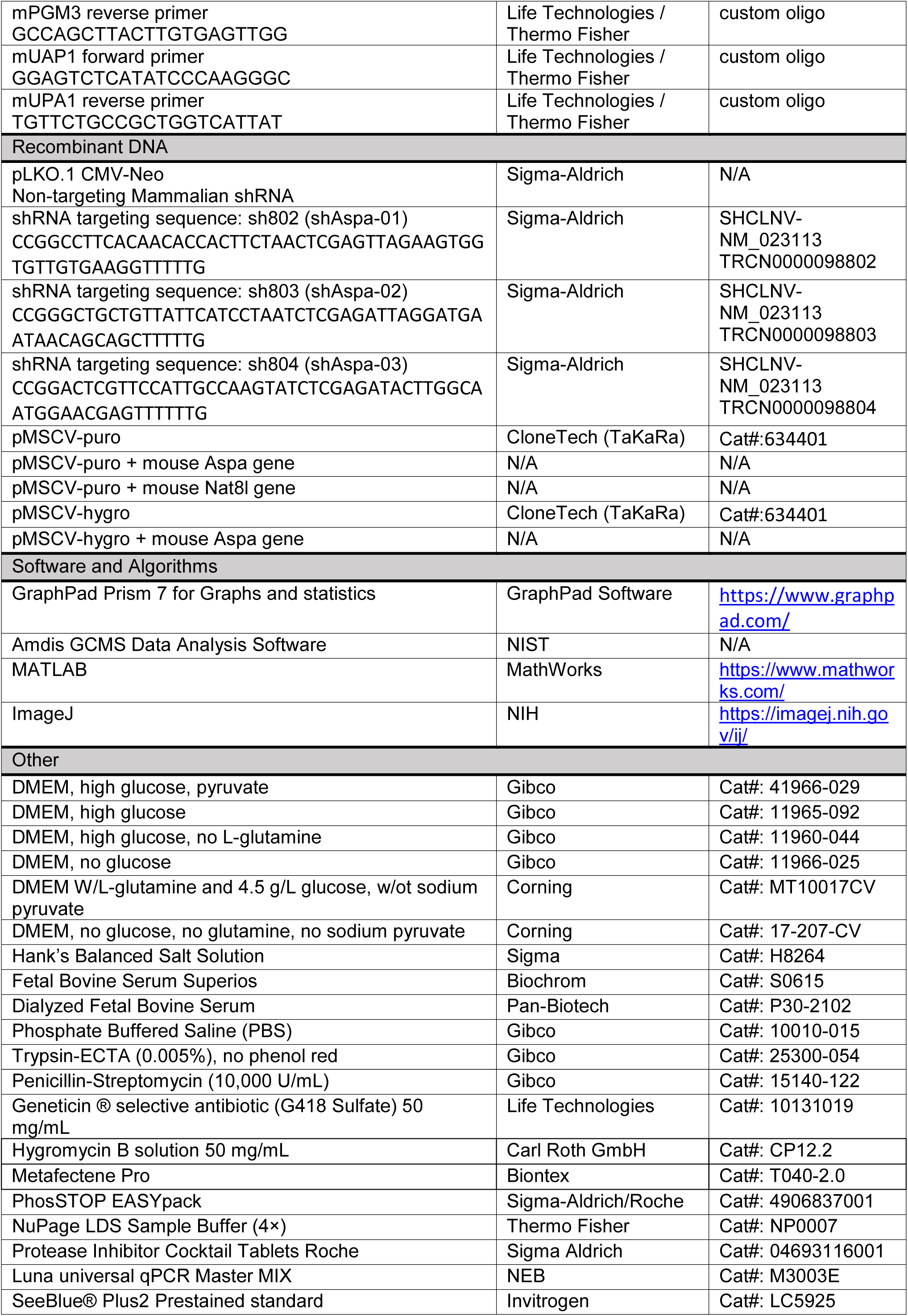
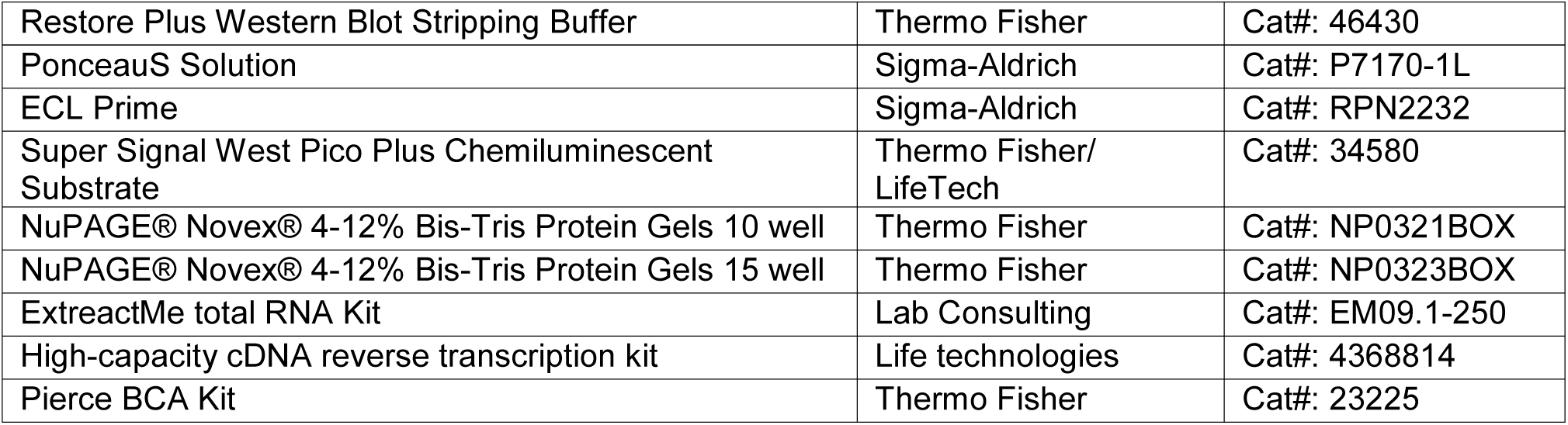

### CONTACT FOR REAGENT AND RESOURCE SHARING

Further information and requests for resources and reagents should be directed to the Lead Contact, Juliane G. Bogner-Strauss

### EXPERIMENTAL MODEL AND SUBJECT DETAILS

#### Cell Lines

All cells were sub-cultured in Dulbecco’s Modified Eagle’s Medium (DMEM, (25 mM Glucose, 4 mM Glutamine, 1 mM Sodium Pyruvate) (GIBCO)) supplemented with 10% fetal bovine serum and 50 units/mL Penicillin/Streptomycin (DMEM++) and incubated at 37°C with 5% CO_2_. All cells utilized tested negative for Mycoplasma.

#### Animals

The study was approved by the institutional ethics committee and experiments were performed according to the guidelines of the Austrian Federal Ministry of Science and Research. 500,000 LLC1 cells were injected into flanks of 7-8 weeks old female C57BL/6 mice (Janvier). Tumor sizes were measured using calipers throughout the study and estimated volumes were calculated by using the formula V= (π/6)^*^(length*width^2^) (<u>Gui et al., 2016</u>). Tumors were harvested and blood was collected from anesthetized mice by cervical dislocation performed after 16 days post-injection.

## METHOD DETAILS

### Overexpression of Nat8l and Aspa

To achieve the stable overexpression of Nat8l (Nat8l o/e) or Aspa (Aspa o/e), recombinant retroviral particles expressing Nat8l or Aspa were used. Recombinant retroviruses expressing Nat8l or Aspa were generated by adding 4 µg Metafectene Pro (Biontex Laboratories GmbH) to Phoenix packaging cells at a confluency as great as 75% (cultured in DMEM++ in 5% CO_2_ at 37°C) with either 5 µg of the vector pMSCVpuro-Nat8l (Nat8l o/e) or pMSCVhygro-Aspa or pMSCVpuro-Aspa (Aspa o/e) or the corresponding controls pMSCVpuro (PuroN) for Nat8l or pMSCVhygro (HygA) or pMSCVpuro (PuroA) for Aspa, respectively. Viral particle-containing supernatants were collected 48 hours after transfection and then were filtered (0.45 µm) and stored as 1 mL aliquots at −80°C until use. 24 hours before the virus treatment, 3500 of mouse LLC1 or 10,000 of human A549 or H1299 cancer cells were seeded into 6-well plates with 1 mL of fresh DMEM++ media. The cells were allowed to adhere overnight. One well was prepared as death control where no virus was applied. The next day, a mixture of 1 mL viral supernatants containing 8 µg/mL polybrene and 1 mL fresh media was added to the cells, followed by 48 hours of incubation. Afterward, an additional 1 mL of fresh virus-containing media was added to the cells for one more day. Then, cells were transferred to T75 flasks with fresh media. The following days, cells were selected with 2.5 µg/mL puromycin or 500 µg/mL hygromycin for 6-7 days. If necessary, cells were split into new flasks.

### Transient overexpression of Aspa in Nat8l o/e cells

In order to investigate effects of transient overexpression of murine Aspa on top of Nat8l o/e on proliferation, 10,000 LLC1 Nat8l o/e cells were seeded to 6-well plates containing 1.5 mL fresh media. Furthermore, mixtures of 0.5 µg pcDNA HisMax C vectors with or without Aspa and 2 µL Metafectane Pro from Biontex in 100 µL serum free DMEM were prepared and added to the cells. After 24 hours, the cells were used for proliferation assays and for the verification of transient Aspa expression via mRNA.

### Aspa-silencing using short-hairpin RNA (shRNA) in LLC1 cells

Non-target shRNA (NTC) and shRNA lentivirus targeting Aspa (shAspa-x; sh802, sh803, sh804) were purchased from Sigma (MISSION™ shRNA lentiviral particles). Prior to transduction, 10,000 LLC1 cells were seeded to 6-well plates to allow the cells to adhere overnight. The next day, media was changed and lentiviral particles with a multiplicity of infection (MOI) of 6 with 8 µg/mL polybrene in DMEM++ were added to the cells. After 4 days, the cells were passaged to T25 flasks and selected with 600 µg/mL neomycin (G-418) for the following 6-7 days.

### Proliferation/Survival experiments

10,000 cells/well were seeded into 6-well plates one day prior to the start of the treatment, allowing the cells to adhere overnight. The next day, cells of the reference plate were washed, trypsinized, re-suspended in PBS and counted with an automated cell counter (Bio Rad) to determine initial cell number. Next, the experimental wells were washed once very carefully with 1 mL PBS, followed by the application of 2 mL of the conditioned media. The cells were treated for up to 2-4 days. The final cell number was determined after trypsinization using the automated cell counter (Bio Rad). The doubling/day rate was calculated by the following formula: Doubling/day = LOG_2_(Final cell count/Initial cell count)/Day

Various conditioned media were used to investigate the effects of nutrients on cancer cell proliferation: DMEM (25 mM Glucose, 4 mM Glutamine, 1 mM Sodium Pyruvate supplemented with 10% dialyzed fetal bovine serum and 50 units/mL Penicillin/Streptomycin), DMEM without pyruvate (-PYR, mainly referred as DP throughout the main text), DMEM without glutamine/pyruvate (-GLN) or DMEM without glucose/pyruvate (-GLC). DP media was used as the control media condition, if not stated otherwise.

For the glucose starvation experiments, two different conditioned media were used. Mouse cancer cell lines were treated either with zero glucose media (0 GLC; composition: Dulbecco’s Modified Eagle’s Medium (DMEM) containing 0 mM glucose, 4 mM glutamine supplemented with 10% dialyzed fetal bovine serum (FBS) and 50 units/mL penicillin/streptomycin (P/S)) or media supplied with 5 mM 2-deoxyglucose (2-DG; composition: Dulbecco’s Modified Eagle’s Medium (DMEM) containing 25 mM glucose, 4 mM glutamine supplemented with 10% dialyzed fetal bovine serum (FBS), 50 units/mL penicillin/streptomycin and 5 mM 2-deoxyglucose). Human cancer cell lines were treated with either 0 GLC media or media supplied with 10 mM 2-DG (composition: Dulbecco’s Modified Eagle’s Medium (DMEM) containing 25 mM glucose, 4 mM glutamine supplemented with 10% dialyzed fetal bovine serum (FBS), 50 units/mL penicillin/streptomycin and 10 mM 2-deoxyglucose). All media were sterile-filtered prior to use.

For the rescue studies, zero glucose (0 GLC) or 2-DG starvation treatments were used and supplemented with additional 10 mM NAA, 10 mM acetate (ACET), 10 mM aspartate (ASP) or the combination of 10 mM acetate and 10 mM aspartate (A/A).

### Western Blot Analysis

Protein was harvested after removing conditioned media and washing carefully with 1 mL of PBS by scraping the cells from the wells with 80 µL RIPA buffer (composition: 50 mM Tris-HCL (pH 8.0), 150 mM NaCl, 1% TritonX-100 or NP-40, 0.5 % Na-deoxycholate, 0.1 % SDS) with 1× Protein Inhibitor Cocktail (PIC) and 1× Phospho Stop (PS). For the protein isolation of tumors, samples were homogenized in 300 µL RIPA buffer and left on ice for up to 20 min. Afterward, the samples were centrifuged for 15 min at 4°C and the maximum speed. Protein concentrations were determined using the BCA protein assay kit from Pierse according to the manufacture’s protocol. Protein lysates (20-40 µg) were loaded to a 4-12 % gradient Bis-Tris gel (NUPAGE, Thermo Fisher Scientific Inc.), which was blotted to a nitrocellulose membrane for 1.5 hours at 500 mA and 4°C. After the transfer, the membrane was first blocked either with 5 % BSA or 5 % dry milk in Tris-Buffered Phosphate Saline (TBST) depending on which antibody was applied in the following steps. The 1^st^ Antibody was applied to the membrane overnight at 4°C. The next day, for chemiluminescent detection, the membrane was exposed to the horseradish peroxidase-conjugated secondary antibody. For detection, SuperSignal West Pico Chemiluminescent Substrate or Amersham ECL Prime was used. GAPDH or PonceauS staining served as the loading controls. For quantification, ImageJ was used.

### Quantitative Real-Time PCR

Total cellular RNA was isolated using the “Total RNA isolation kit” from Sigma, according to the manufacturer’s protocol. To isolate the RNA from the tumor tissue, the TRIzol reagent from Thermo Fisher Scientific Inc. was used. High Capacity cDNA Reverse Transcription kit from Applied Biosystems was used to generate cDNA by reverse transcription. The expression of individual genes was determined by using LUNA (NEB) quantitative Real-Time PCR (qRT-PCR) on the StepOne Plus Real-Time PCR System (Applied Biosystems) in 96-well plates using a 4 pmol forward (fw) and reverse (rv) primer mixture, 4 ng cDNA and 8 µL LUNA per well. The expression values of each gene were normalized to the house keeping gene m36b4 (Rplp0) (for mouse cell lines) or hRPLP0 (for human cancer cell lines) and then normalized to the mean of control group for each gene followed by calculating fold change.

### NMR Spectroscopy

NMR spectroscopy measurements were performed as previously described (Alkan et al, 2018). Methanol, sodium phosphate, dibasic (Na_2_HPO_4_), sodium hydroxide, hydrochloric acid (32 % m/v), and sodium azide (NaN_3_) were obtained from VWR International (Darmstadt, Germany). 3(trimethylsilyl)propionic acid-2,2,3,3-d4 sodium salt (TSP) was obtained from Alfa Aesar (Karlsruhe, Germany). Deuterium oxide (D_2_O) was obtained from Cambridge Isotope laboratories, Inc. (Tewksbury, MA). Deionized water was purified using an inhouse Milli-Q® Advantage Water Purification System from Millipore (Schwalbach, Germany). All chemicals were used with no further purification. The phosphate buffer solution was prepared by dissolving 5.56 g of anhydrous NaH_2_PO_4_, 0.4 g of TSP, and 0.2 g NaN_3_, in 400 ml of deionized water and adjusted to pH 7.4 with 1 M NaOH and HCl. Upon addition of deionized water to a final volume of 500 ml the pH was readjusted to pH 7.4 with 1 M NaOH and HCl. The buffer was lyophilized and taken up in 500 ml D_2_O to obtain NMR buffer in D_2_O.

For sample preparation; sub-confluent LLC1 cells from 150 cm dishes or 20-150 mg tissues collected in PBS. Tissues were lysed using 1.0 mm diameter zirconia beads (Carl Roth) by vigorous shaking in Precellys 24 (Bertin Technologies) for 2×20 seconds and by sonication 3×10 seconds. Cell lysates were lysed solely via sonication. For quenching the metabolites; one volume of cell lysate, cell media, tissue lysate or plasma is mixed with two volume of cold methanol, incubated at −20 °C for at least 1 hour, and centrifuged at 13,000 rpm for 30 min to pellet proteins. Supernatants were transferred to fresh vials and dried for 4 hours at room temperature. 500 μL of NMR buffer in D_2_O were added to the samples, re-dissolved and transferred to 5 mm NMR tubes.

All NMR experiments were performed at 310 K on a Bruker Avance III 500 MHz spectrometer equipped with a TXI probe head. The 1D CPMG (Carr−Purcell−Meiboom−Gill) pulse sequence (cpmgpr1d, 73728 points in F1, 12019.230 Hz spectral width, 1024 transients, recycle delay 4 s), with water suppression using pre-saturation, was used for 1H 1D NMR experiments.

Bruker Topspin version 3.1 was used for NMR data acquisition. The spectra for all samples were automatically processed (exponential line broadening of 0.3 Hz), phased, and referenced to TSP at 0.0 ppm using Bruker Topspin 3.1 software (Bruker GmbH, Rheinstetten, Germany).

Preprocessed 1D NMR spectra were transferred into Matlab R2014a (The Mathworks, Inc., USA) and processed using lab-written scripts. 1D NMR spectra were referenced to the TSP peak at 0 ppm, before peak alignment by recursive segment-wise peak alignment (RSPA). The spectrum with the highest correlation to the other spectra was used as the alignment reference. The spectral data between 0.5 and 10.0 ppm were extracted for multivariate analysis, after removal of the residual water (4.5 – 5.0 ppm) and TSP (−0.5 – 0.5 ppm) regions. Prior to multivariate analysis, the spectra were normalized by probabilistic quotient normalization (PQN). Multivariate analysis was performed on mean-centered data by orthogonalized partial least squares discriminant analysis (OPLS-DA) validated by leave-one-subject-out cross validation and permutation testing (n = 1000, significance for pperm ≤ 0.05). Metabolite reference chemical shifts were taken from the Madison-Qingdao Metabolomics Consortium Database (http://mmcd.nmrfam.wisc.edu/) database and metabolites were cross-checked using reference compounds if necessary.

### Isotope tracing and gas chromatography - mass spectroscopy analysis

For metabolic tracing studies, 150,000 to 200,000 cells/well were seeded in 6-well plates overnight. Cells were washed three times, then 5 mM [U^13^C]glucose or 4 mM [U^13^C]glutamine (Cambridge Isotope Laboratories) containing DMEM (10% dialyzed serum, no pyruvate) was applied. After 24 hours of culture, metabolites were extracted from cells or media (10 µL) in 80 % methanol in water containing 1 µg/sample norvaline and then were dried under nitrogen gas. The polar metabolites were derivatives and thus were measured as described previously (Lewis et al., 2014). Relative metabolite abundances were calculated by integrating ion peak area and normalizing to norvaline, then altering based on the cell numbers from identical plates. Mass isotopomer distributions of each ion peak were determined after natural abundance corrections adapted from: Fernandez et al., 1996.

### Quantification and Statistical Analysis

If not stated otherwise, all shown results are mean values +/- standard deviation (SD) of at least three independent experiments, or a data set shows one representative experiment of at least three independent experiments. Statistical significance was calculated via unpaired two-tailed students t-test or two-way ANOVA tests using GraphPad Prism Software. Significance levels: * p ≤ 0.05, ** p ≤ 0.01, *** p ≤ 0.001.

