## Supplementary material for "N-acetylaspartate improves cell survival when glucose is limiting": Suplemental Figures

### Supplemental Information

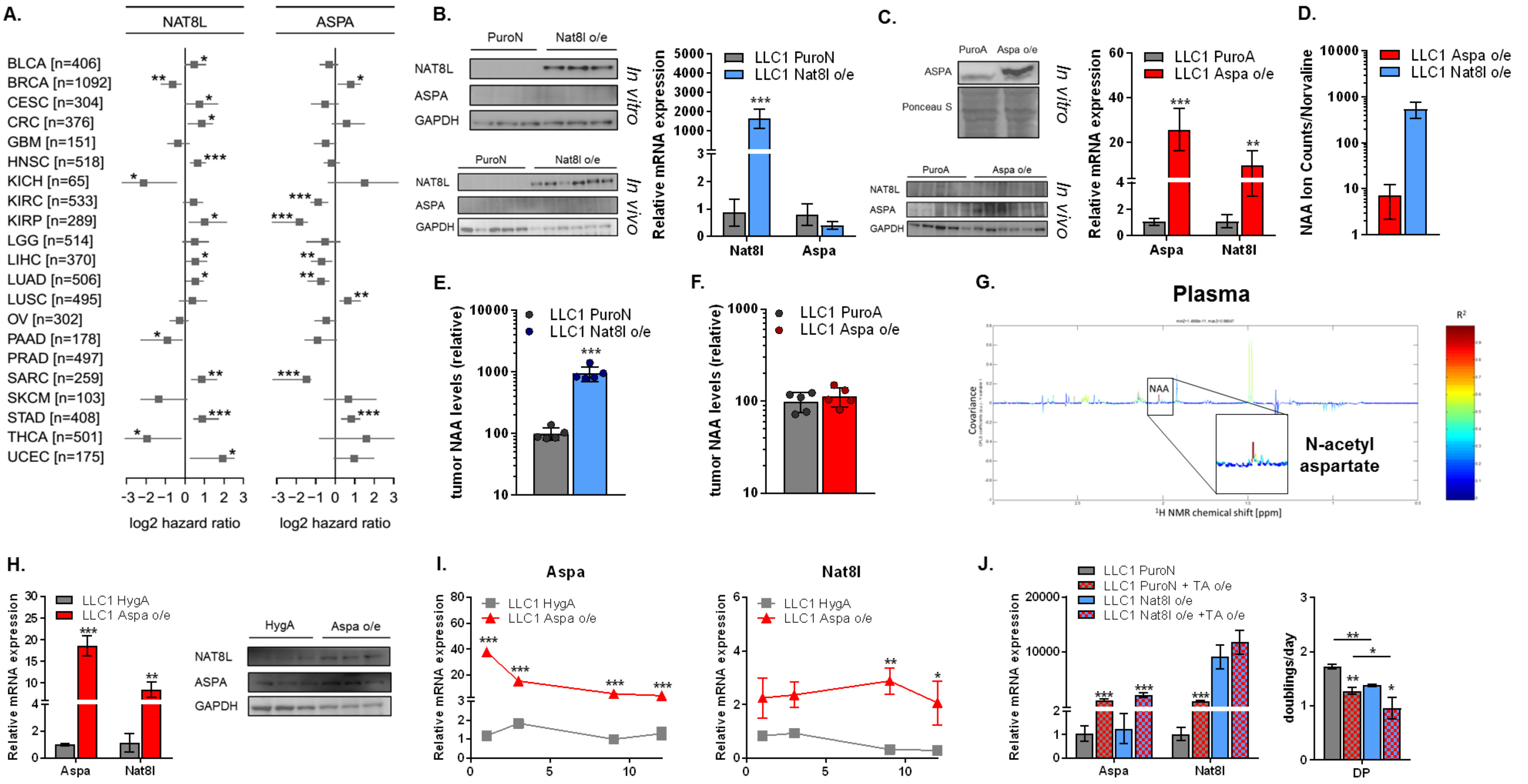


**Figure S1. Impacts of Nat8l and Aspa overexpressions on patient survival, tumor growth and cell proliferation.**

**(A)** Correlation of NAT8L or ASPA mRNA expression with better (negative log2hazard ratio) or poor (positive log2hazard ratio) survival of patients from various cancers. Lower log2hazard ratio for a gene means the group of the patients with tumors with lower expression that gene has worse overall survival than the higher expressing group. Data is adopted from the TCGA database. Patients with higher or lower NAT8L or ASPA mRNA expression was divided into two groups with a cut-off line where the separation of these groups was significantly most meaningful. **(B)** Nat8l and Aspa protein expressions in control (PuroN) and Nat8l overexpressing (Nat8l o/e) LLC1 (top, left) cells (*in vitro*) or (bottom, left) tumors (*in vivo)* using Western Blot analysis. (right) mRNA expressions of Nat8l and Aspa in LLC1 control and Nat8l o/e tumors determined via quantitative-Real-Time PCR analysis. **(C)** Nat8l and Aspa protein expressions in control (PuroA) and Aspa overexpressing (Aspa o/e) LLC1 (top, left) cells (*in vitro*) or (bottom, left) tumors (*in vivo)* using Western Blot analysis. (right) mRNA expressions of Nat8l and Aspa in LLC1 control and Aspa o/e tumors determined via quantitative-Real-Time PCR analysis. **(D)** Ion counts of N-acetylaspartate (NAA) in Aspa and Nat8l overexpresing cells, normalized to norvaline and cell number**. (E, F)** Relative NAA levels in control (PuroN, PuroA), (E) Nat8l overexpressing (Nat8l o/e) and (F) Aspa overexpressing (Aspa o/e) LLC1 tumors, determined via 1H-NMR sprectroscopy (n=5). **(G)** $^{1}$H-NMR spectroscopy of plasma samples of mice with Nat8l overexpressing tumors, normalized to the mice with control tumors. Positive covarience (elevated peaks) of a relevant metabolite indicate an increase in Nat8l tumors compared to controls. Higher $R^2$ values (red) indicate significant changes (n=3). **(H)** Verification of Aspa overexpressions in LLC1 cells using (left) qPCR and (right) Western Blot analysis. **(I)** Relative changes in mRNA experssions of Aspa and Nat8l in control (HygA) and Aspa overexpressing (Aspa o/e) cells over 12 days. **(J)** Proliferation rate of control (PuroN) and Nat8l o/e LLC1 cells without or with additional transient Aspa o/e (TA o/e) cultured in pyruvate-free DMEM (DP) (n=3). (left) mRNA expression levels of Aspa and Nat8l in LLC1 control (PuroN) and Nat8l o/e cells without or with transient Aspa o/e (TA o/e) (n=3).

All figures denote mean ± SDs. Significance levels: * $p \le 0.05$, ** $p \le 0.01$, *** $p \le 0.001$

**(related to Figure1)**

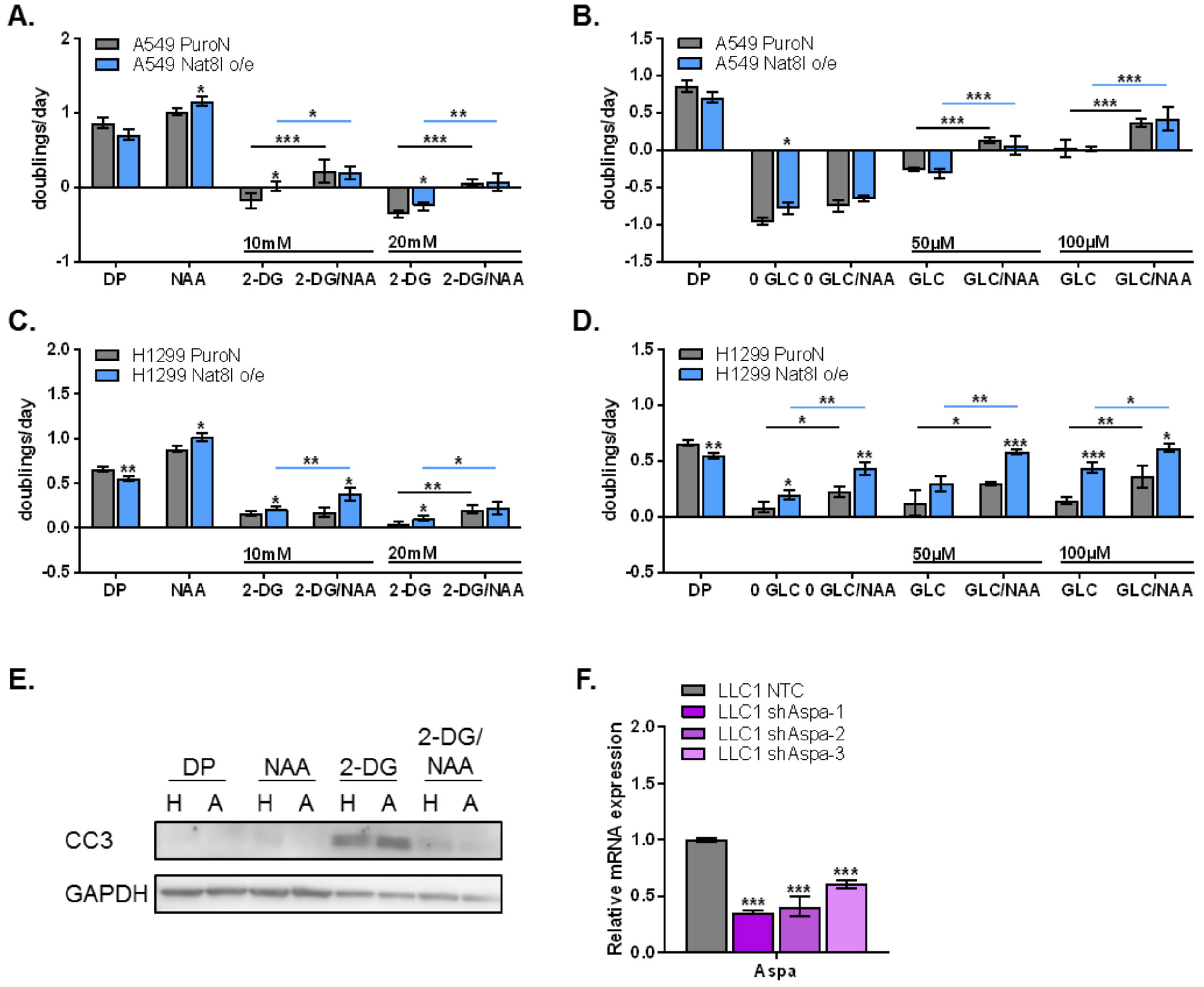


**Figure S2. NAA improves cell proliferation and survival in low glucose.**

**(A-D)** Proliferation/survival rate (shown as doublings/day) of control (PuroN) and Nat8l overexpressing (Nat8l o/e) A549 and H1299 cells in DMEM without pyruvate (DP) in the absence or presence of 10 mM or 20 mM 2-deoxyglucose (2-DG), 0 µM (0 GLC), 50 µM, or 100 µM glucose or 10 mM N-acetylaspartate (NAA) for 96 hours (n=3). All conditions contain 10% dialysed FBS. **(E)** Western Blot analysis of Cleaved Caspase 3 (CC3) protein in control (H) and Aspa overexpressing (A) LLC1 cells in the presence and absence of 5 mM 2-deoxyglucose (2-DG) and 10 mM N-acetylaspartate (NAA). **(F)** Verification of Aspa expression in control (NTC) and Aspa-knockdown (shAspa) LLC1 cells using qPCR analysis.

All figures denote mean ± SDs. Significance levels: * $p \le 0.05$, ** $p \le 0.01$, *** $p \le 0.001$

**(related to Figure 2)**

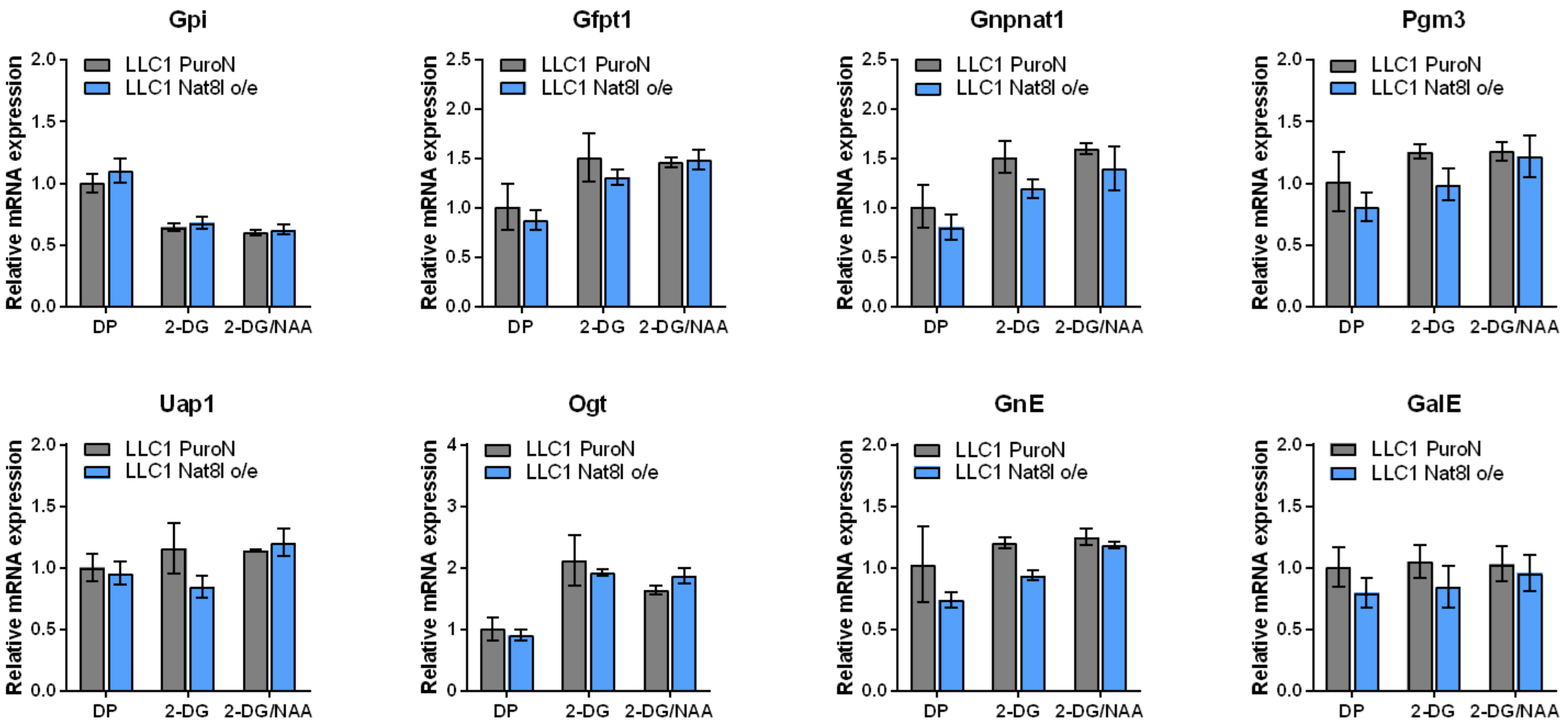


**Figure S3. NAA or Nat8l overexpression do not regulate hexoseamine biosynthesis pathway on transcription level.**

mRNA expression of Gpi, Gfpt1, Gnpnat1, Pgm3, Uap1, Ogt, GnE and GalE in control and Nat8l o/e LLC1 cells cultured in pyruvate-free DMEM (DP) and treated with either 5 mM 2-deoxyglucose (2-DG) or 5 mM 2-DG and 10 mM NAA (2-DG/NAA). Gene name abbreviations: Glucose-6-Phosphate Isomerase (Gpi); Glutamine--Fructose-6-Phosphate Transaminase 1 (Gfpt1); Glucosamine-Phosphate N-Acetyltransferase 1 (Gnpnat1); Phosphoglucomutase (Pgm3); UDP-N-Acetylglucosamine Pyrophosphorylase 1 (Uap1); O-Linked N-Acetylglucosamine (GlcNAc) Transferase (Ogt); Glucosamine (UDP-N-Acetyl)-2-Epimerase/N-Acetylmannosamine Kinase (GnE); UDP-Galactose-4-Epimerase (GalE)

**(related to figure 3)**

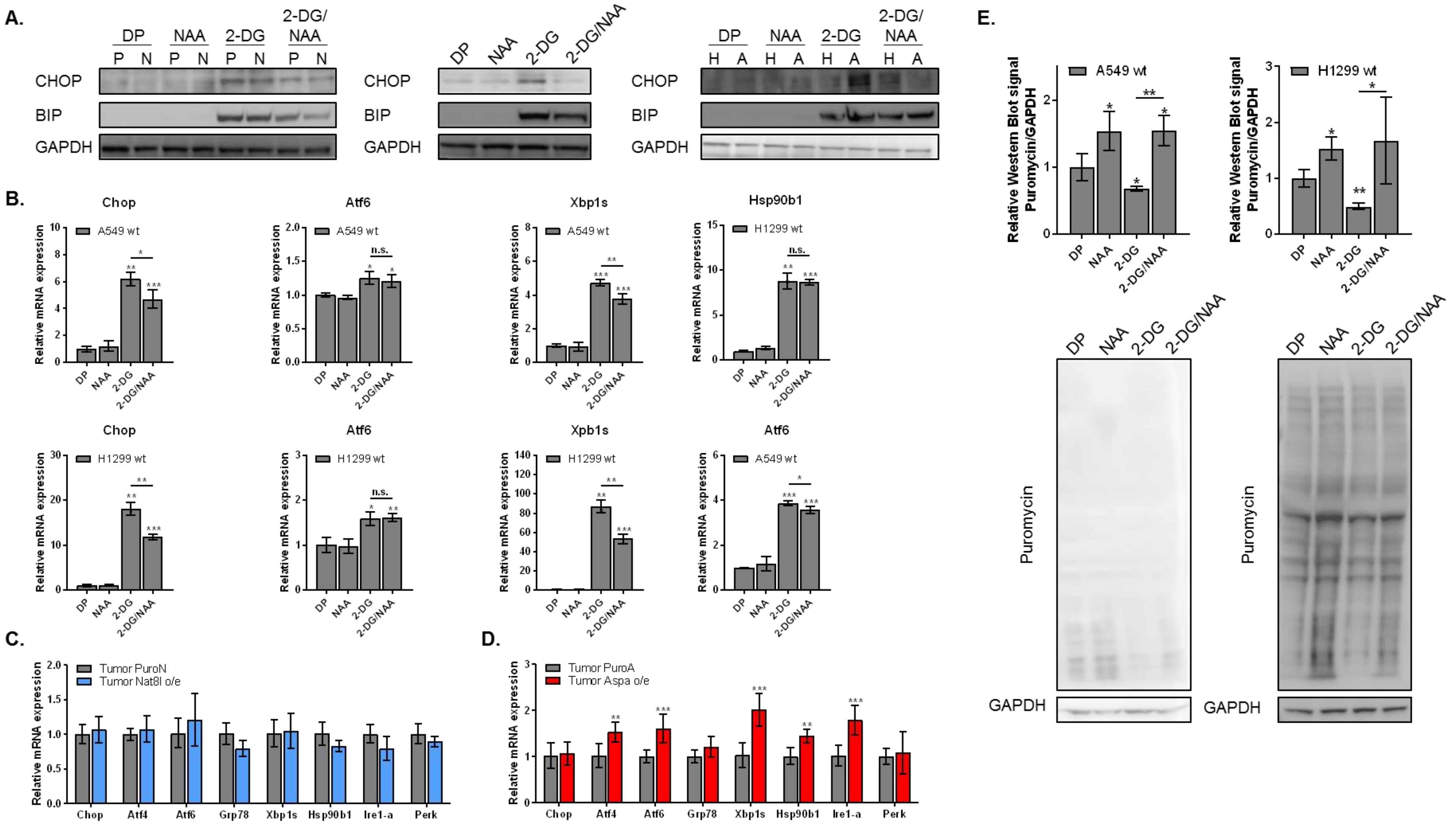


**Figure S4. NAA improves ER stress response and protein synthesis when glucose metabolism is inhibited.**

**(A)** CHOP and BIP protein expression in Nat8l o/e (left), wild-type (middle) and Aspa o/e (right) cells, assessed using Western blot analysis. Cells were cultured in pyruvate-free DMEM (DP) and treated with 10 mM N-acetylaspartate (NAA) in either the presence or absence of 5 mM 2-deoxyglucose (2-DG) (n=3). GAPDH was used for normalization. One representative replicate is shown of 3. Mean +/- standard deviations are shown. Abbreviations: P: control (PuroN), N: Nat8l o/e, H: control (HygA), A:Aspa o/e. **(B)** Relative mRNA levels of Chop, Xbp1s, Atf6, and Hsp90b1 in (top) A549 and (bottom) H1299 cells in the presence and absence of 10 mM 2-deoxyglucose (2-DG) and 10 mM N-acetylaspartate (NAA). Rplp0 is used as reference gene (n=3). **(C-D)** mRNA expression of Chop, Atf4, Atf6, Xbp1s, Grp78, Ire1-alpha, Hsp90b1 and Perk in control PuroN or PuroA and (C) Nat8l o/e or (D) Aspa o/e tumors respectively (n=5). Mean +/- standard deviations are shown. Gene name abbreviations: DNA Damage Inducible Transcript 3 (Chop), Activating Transcription Factor 4 (Atf4), Activating Transcription Factor 6(Atf6), spliced X-Box Binding Protein 1 (Xbp1s), Heat Shock Protein Family A (Hsp70) Member 5 (Grp78), Endoplasmic Reticulum To Nucleus Signaling 1 (Ire1-alpha), Heat Shock Protein 90 Beta Family Member 1 (Hsp90b1), Eukaryotic Translation Initiation Factor 2 Alpha Kinase 3 (Perk) **(E)** Western blot analysis of the puromycin incorporation into protein synthesis in A549 and H1299 cells after 15 minutes of 90 µM puromycin treatment in the presence and absence of 10 mM 2-deoxyglucose (2-DG) and 10 mM N-acetylaspartate (NAA). **(related to figure 4)**
